# *Large1* Gene Transfer in Older *myd* Mice with Severe Muscular Dystrophy Restores Muscle Function and Greatly Improves Survival

**DOI:** 10.1101/2021.10.28.466309

**Authors:** Takahiro Yonekawa, Adam J. Rauckhorst, Sara El-Hattab, Marco A. Cuellar, David Venzke, Mary E. Anderson, Hidehiko Okuma, Alvin D. Pewa, Eric B. Taylor, Kevin P. Campbell

**Author notes:** Correspondence and requests for materials should be addressed to K.P.C. **Author Information** The authors declare no competing financial interests.

## Abstract

Muscular dystrophy is a progressive and ultimately lethal neuromuscular disease due to lack of therapeutic options that restore muscle function. Gene editing and gene transfer hold great promise as therapies for various neuromuscular diseases when administered prior to the onset of severe clinical symptoms. However, the efficacy of these strategies for restoring neuromuscular function and improving survival in the late stages of muscular dystrophy with severe muscle pathophysiology is unknown. Dystroglycanopathies are muscular dystrophies characterized by extensive skeletal muscle degeneration and, in many cases, are accompanied by eye and brain abnormalities. Thus far, mutations in at least eighteen human genes are known to cause dystroglycanopathies, including those in the *like-acetylglucosaminyltransferase-1* (*LARGE1*) gene. *LARGE1* encodes a xylosyl- and glucuronosyltransferase that modifies α-dystroglycan (α-DG) with matriglycan, a linear repeating disaccharide of alternating xylose and glucuronic acid that binds to the laminin G-like domains of extracellular matrix proteins with high affinity. *Large*^*myd*^*/Large*^*myd*^ (*myd)* mice lack expression of *Large1*, and exhibit severe skeletal muscle pathophysiology, impaired mobility, and a drastically reduced lifespan (50% survivorship at 35 weeks of age). Here, we show that systemic delivery of AAV2/9 CMV *Large1* (AAV*Large1*) in >34-week-old *myd* mice with advanced disease restores matriglycan expression, attenuates skeletal muscle pathophysiology, improves motor and respiratory function, and normalizes systemic metabolism, which collectively and dramatically extends survival. Our results demonstrate that in a mouse model of muscular dystrophy, skeletal muscle function can be restored, illustrating its remarkable plasticity, and that survival can be greatly improved even after the onset of severe skeletal muscle pathophysiology.

## INTRODUCTION

Substantial progress has been made toward unraveling the genetic basis of various muscular dystrophies, yet existing therapies are limited and do not prevent the unchecked skeletal-muscle degeneration that eventually leads to death. Gene therapy is an attractive approach for these individuals as it enables the delivery of a functional copy of a gene or repairs the mutated locus (for reviews see 1-5). Recently, AAV-mediated gene transfer has shown great promise for treating patients with neuromuscular diseases, such as spinal muscular atrophy, when administered before severe clinical symptoms arise (6). Similarly, preclinical studies in which adenoviral gene transfer or gene editing is administered at a young age prior to the onset of severe symptoms shows great promise in various mouse models of muscular dystrophy (7-15). However, the efficacy of AAV-mediated gene therapy in treating severe muscle degeneration associated with advanced stages of muscular dystrophy is unknown.

Dystroglycanopathies are a group of congenital/limb-girdle muscular dystrophies with or without eye and brain abnormalities and are caused by defects in *O*-glycosylation or post-translational processing of α-dystroglycan (α-DG), the cell-surface subunit of dystroglycan (DG) (16-18). DG is a widely expressed high-affinity extracellular matrix (ECM) receptor (19) that is highly glycosylated and involved in a variety of physiological processes, which include maintaining the integrity of the skeletal muscle membrane and the structure and function of the central nervous system (16-17). In skeletal muscle, DG is part of the dystrophin-glycoprotein complex, which establishes a continuous link between laminin-G-like (LG) domains of ECM-resident proteins (laminin, agrin, and perlecan) and the cytoskeleton (19-22). Dystroglycanopathies are characterized by an absence or reduction in matriglycan, a unique heteropolysaccharide on α-DG that binds with high-affinity to laminin G-like domains of ECM proteins (23-24). Thus far, at least eighteen causative genes for dystroglycanopathies have been identified in humans, including mutations in the *like-acetylglucosaminyltransferase-1* (*LARGE1)* gene that cause a severe form of congenital muscular dystrophy (25-27).

*Large*^*myd*^*/Large*^*myd*^ (*myd)* mice are an excellent model for therapeutic studies of dystroglycanopathies because they lack matriglycan and exhibit severe skeletal muscle pathophysiology, impaired mobility, reduced body weight, and a drastically reduced lifespan (16, 28-30). Using both non-invasive and invasive techniques, we performed a comprehensive study to test the ability of systemic AAV-mediated gene transfer of *Large1* to improve skeletal muscle function and survival in *myd* mice with advanced disease. We show that this treatment in older *myd* mice with severe muscular dystrophy attenuates skeletal muscle degeneration and improves motor and respiratory function as well as systemic metabolism, which collectively extends survival of *myd* mice. Thus, our study provides proof-of-concept that functional and pathological features of muscular dystrophy are treatable, even after severe muscle degeneration is established.

## RESULTS

To evaluate the therapeutic effect of treating established dystroglycanopathy, we targeted aged *myd* mice as they exhibited severe muscle pathophysiology. Prior to gene transfer studies, we analyzed the progression of disease in *our Large*^*myd*^*/Large*^*myd*^ (*myd*) mouse colony on the C57BL/6 genetic background (92.5% C57BL/6) from the newborn period to 35 weeks of age. *myd* mice were generated by mating heterozygous +/*myd* mice and were significantly smaller than their littermates (+/+ or +/*myd*) at postnatal day three, weighing 1.8 ± 0.3 g (mean ± SD, n = 18) versus 2.2 ± 0.5 g (n = 143). *myd* mice also showed consistently slower growth compared to littermate controls, although males and females continued to grow until 20 or 15 weeks of age, respectively, after which they exhibited a significant decline in body weight (Figs. S1a and b). At 35 weeks of age, the weight of *myd* mice was markedly less than that of littermate controls: 21.4 ± 2.7 g (*myd*) versus 42.4 ± 6.1 g (littermate controls) in male and 16.7 ± 2.7 (*myd*) versus 34.2 ± 5.5 g (littermate controls) in female. The survival of *myd* mice was considerably shorter than that of littermate controls, with only 35% surviving to 40 weeks of age and no mice living past 60 weeks of age (Fig. S1c). Additionally, thoracic kyphosis and muscle wasting were evident at ∼40 weeks of age in *myd* mice (Fig. S1d). Fibrous and adipose tissue infiltration was prominent in *myd* mice that were older than 35 weeks, with numerous small, round-shaped fibers observed upon histological analysis, indicative of end-stage pathology (Figs. S2a, S2b). Matriglycan positive α-DG (functionally glycosylated) was localized in the muscle sarcolemma of C57BL/6 (WT) mice but absent in *myd* mice (Fig. S2c).

We next performed systemic AAV-mediated gene transfer of *Large1* in adult *myd* mice (at approximately 50% survivorship age) with severe skeletal muscle pathophysiology and assessed the effects on muscle function and survival. We randomly assigned 35 *myd* mice that were 34 weeks or older to untreated or treated groups. Twenty-one mice were left untreated whereas fourteen mice were injected once with AAV*Large1* at age 38.4 ± 2.8 weeks (mean ± SD). A series of non-invasive and invasive therapeutic readouts were compared between the two groups, including: body weight, survival, muscle and respiratory function, muscle physiology, histological evaluation, plasma metabolomics, and biochemical analysis of α-DG. Mice were weighed weekly and locomotor activity was assessed every four weeks (see Fig. 1a experimental outline). Body weight and locomotor activity were compared between baseline (i.e., at enrollment for untreated group or before AAV*Large1* injection for treated group) and each assessment time point. Respiratory function was assessed before untreated mice were euthanized or when AAV-treated mice reached 60 weeks of age. Muscle physiology was evaluated immediately prior to euthanasia and tissues were subsequently harvested for further analysis.

**Figure 1.**
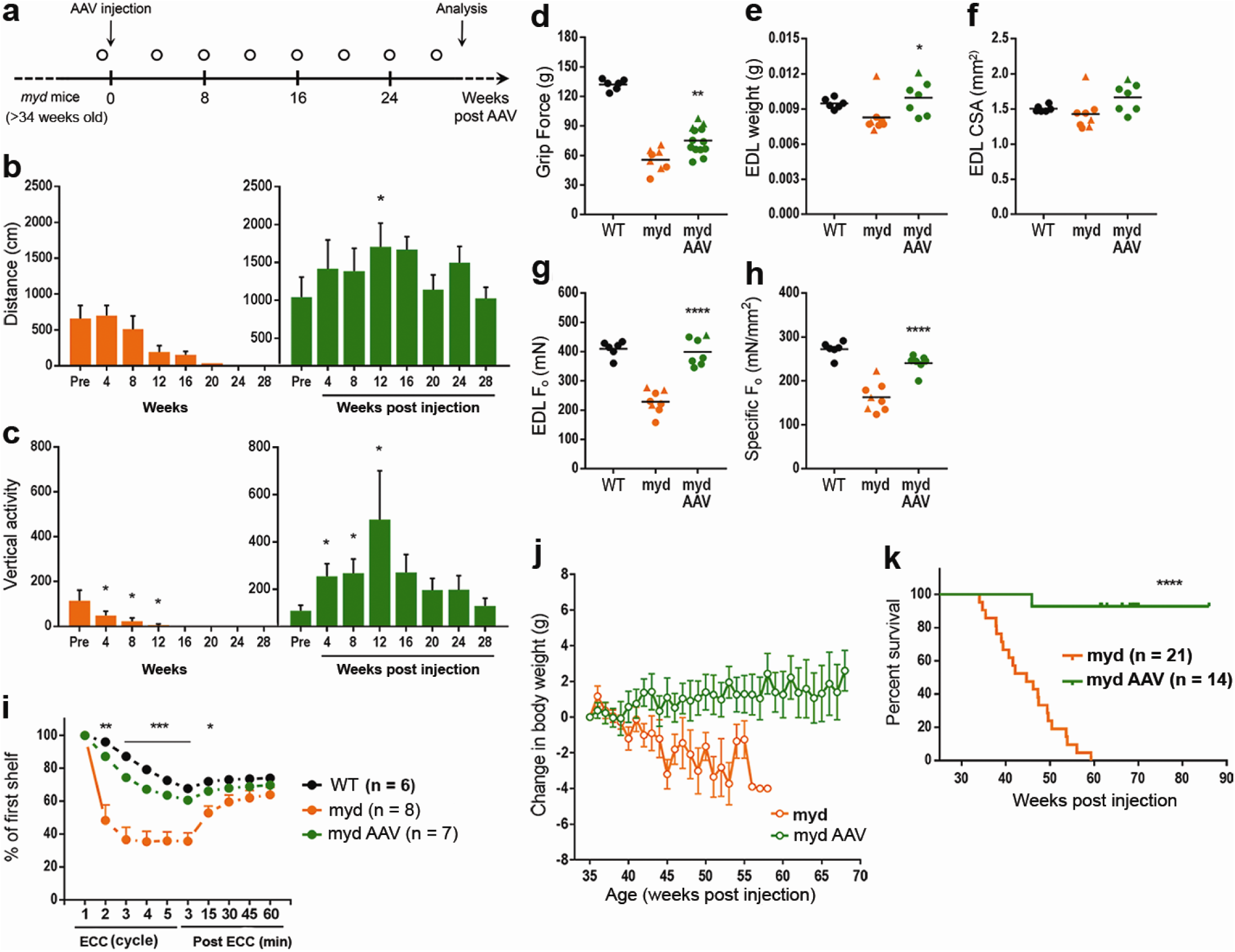
*Large1* gene transfer improves skeletal muscle function and extends survival. **a**, Experimental outline. Mice were weighed weekly; open circle, locomotor activity determined. **b, c** Spontaneous locomotion (distance) and rearing (vertical activity). Untreated, orange; treated, green. **d-h**, Forelimb grip strength **d**, EDL muscle weight **e**, EDL muscle CSA **f**, EDL isometric tetanic force (F_o_), **g**, specific isometric tetanic force (F_o_/CSA) **h**, (black; C57BL/6J) (WT)); treated *myd* (myd AAV), age > 60 weeks, untreated *myd* (myd), age 46.5 ± 6.5 weeks. **i**, Percentages of F_o_ in mice at indicated time after eccentric contraction (ECC), relative to F_o_ at 1^st^ ECC. **j**, Changes in body weight relative to weight at 35 weeks of age. **k**, Survival curves of mice in the indicated treated groups. Symbols, individual mice; bars, means ± SEM. C57, black; myd, orange; myd AAV, green.

Untreated *myd* mice (34–35 weeks old) showed a marked reduction in locomotor activity, as measured by distance traveled, or rearing behavior, as measured by vertical activity, and displayed progressive deterioration in motor performance until euthanasia was necessary (Figs. 1b, c) compared to 37-week-old WT mice (Fig. S3). In contrast, AAV-mediated gene transfer of *Large1* in *myd* mice restored locomotor and rearing activities, and mice were able to maintain motor performance until they were euthanized for analysis at age 69.7 ± 5.1 weeks (Figs. 1b, c).

Forelimb grip strength was also markedly reduced in 46.5-week-old *myd* mice relative to WT mice aged 61.1 weeks (Fig. 1d). The muscle weight and cross-sectional area (CSA) of dissected extensor digitorum longus (EDL) muscles were smaller (Figs. 1e, 1f), and the absolute isometric tetanic force (F_o_) and the size-normalized isometric tetanic force (F_o_/CSA) were markedly lower in *myd* mice compared to those of WT mice (Figs. 1g, 1h), suggesting that the muscles were not only atrophic but also highly degenerated. In contrast, AAV*Large1-*treated *myd* mice showed a significant increase in forelimb grip strength (Fig. 1d); however, the muscle weight and CSA of isolated EDL muscles only increased modestly (Figs. 1e, 1f).

Systemic transduction of *myd* mice with AAV*Large1* of *myd* mice also resulted in a significant increase in isometric tetanic force (F_o_) and specific isometric tetanic force (F_o_/CSA) of EDL muscles compared to untreated mice (Figs. 1g, h), demonstrating that even at advanced stages of disease muscle contractile properties can be improved by delivering a functional copy of *Large1*. Additionally, EDL muscles in untreated mice were highly susceptible to injury induced by eccentric contractions (ECCs), and tetanic F_o_ force dropped by 63% after the 3^rd^ ECC (Fig 1i). In contrast, EDL muscles in AAV*Large1-*treated mice tolerated a sequence of five ECCs, although the force drops induced by the ECC protocol were slightly greater than those in WT muscles (Fig. 1i). It should be noted that muscle size, contractile properties, and lengthening contraction-induced muscle damage is similar in EDL muscles from +/+ or +/*myd* mice (Fig. S4).

At 35 weeks of age (just before AAV treatment), there was no significant difference in body weights between *myd* mice that were untreated or treated with AAV*Large1* (Fig. S5a). However, after 35 weeks, untreated *myd* mice displayed a significant decline in body weight that progressed until euthanasia was required (Fig. 1j), whereas systemic gene transfer of *Large1* resulted in a mild gain in body weight (Fig. 1j). All surviving *myd* mice older than 35 weeks of age were very thin, suggesting they had already developed severe muscle atrophy and degeneration, and they continued to decline without treatment (Fig. S5b, left). Indeed, the sizes of the gastrocnemius and quadriceps femoris muscles were markedly smaller in *myd* mice compared to those of WT mice. Nevertheless, AAV-mediated gene transfer of *Large1* resulted in mild growth of these muscles (Figs. S5c–f), although treated mice still looked thin (Fig. S5b, right), suggesting these muscles may be too severely affected at later stages of disease to recover to normal weights. Strikingly, AAV-mediated gene transfer significantly extended the survival of *myd* mice, as all but one mouse injected with AAV*Large1* lived to >65 weeks of age, whereas only 50% of the *myd* mice survived to 45 weeks of age (Fig. 1k).

As expected, *myd* muscles did not express *Large1* and the level of *Large1* expression in +/*myd* muscles was approximately half that observed in muscles from WT or +/+ mice (Fig. 2a). Western blot analysis of WGA-enriched *myd* muscle homogenates demonstrated that the transmembrane subunit of DG, β-dystroglycan (β-DG), was normally expressed whereas functional α-DG was absent in *myd* muscles, as assessed by reactivity to the IIH6 antibody (anti-matriglycan) or the laminin overlay assay (Fig. 2b). In addition, α-DG–ligand binding was severely affected in *myd* muscles compared to WT muscles (Fig. 2c) corresponding to the fact that *myd* EDL muscles were highly susceptible to lengthening contractions (Fig. 1i). Of note, the quadriceps and EDL muscles in +/*myd* mice exhibited no difference in biochemical and physiological features, respectively, when compared to WT or +/+ muscles (Figs. 2b and S4). The level of *Large1* expression in +/*myd* (+/-) muscles was around half of that of WT or +/+ levels (Fig. 2a), and +/*myd* muscles showed no biochemical or physiological abnormalities (Figs. 2b and S4).

**Figure 2.**
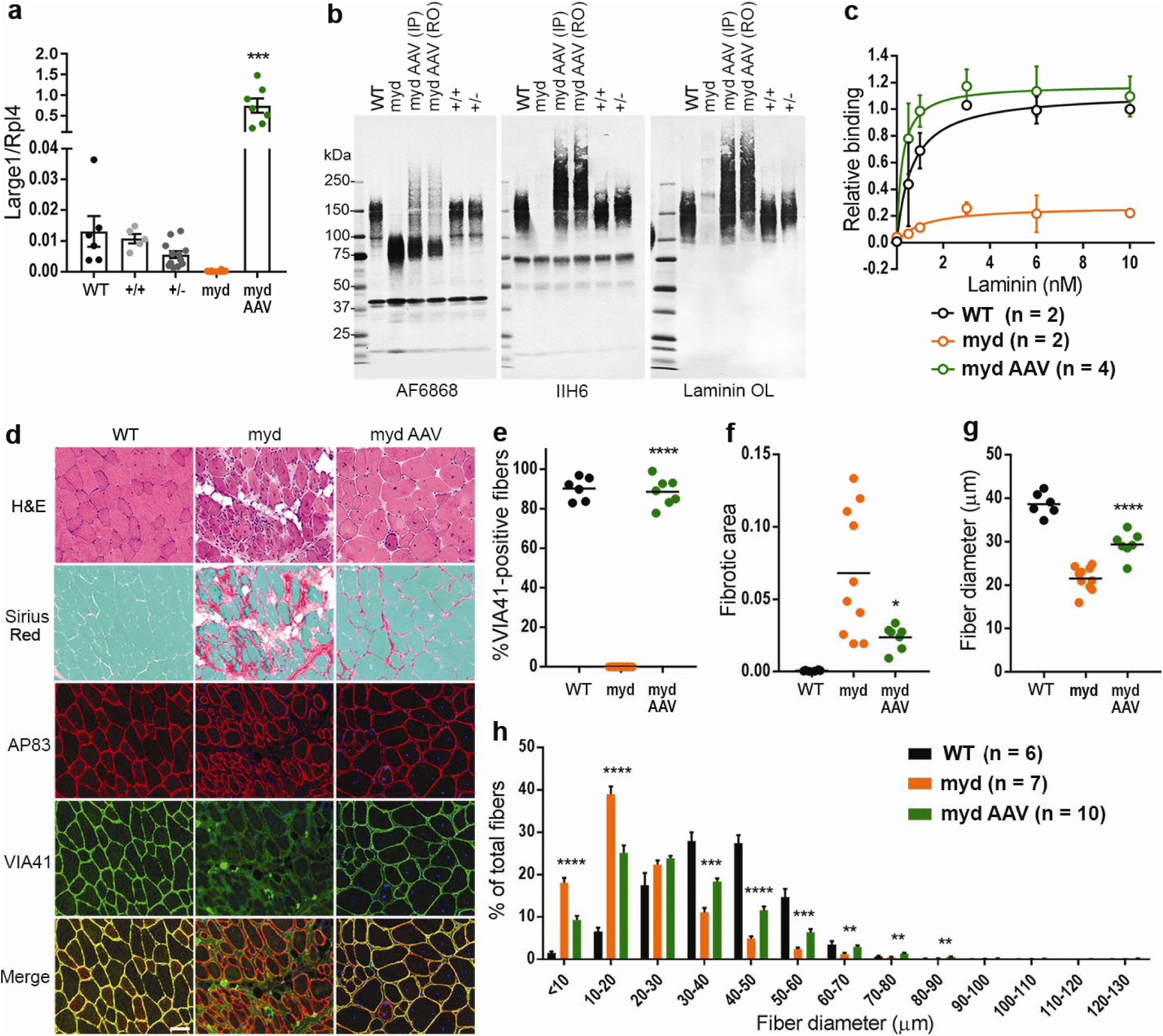
AAV*Large1* restores matriglycan on α-DG and laminin binding in skeletal muscles, which alleviates muscle pathology in *myd* mice. **a**, *Large1* expression. **b**, Western blot of core α-DG and β-DG (AF6868), matriglycan-positive α-DG (IIH6), or laminin (laminin OL) in a. **c**, Solid-phase binding assay. **d**, Representative cryosections stained with H&E or Sirius Red & Fast Green, or used for immunofluorescence: β-DG (AP83); matriglycan-positive α-DG (VIA41). Scale bar: 50μm. **e-h**, Quantitative analysis of sections in d. **e**, VIA41-positive fibers. **f**, Connective tissue deposition. **g**, Average Feret’s diameter of fibers. **h**, Percentage of fibers of indicated diameter. Symbols, individual mice; bars, means ± SEM. WT, C57Bl/6J; +/+, *Large*^*+/+*^; +/-, *Large*^*+/-*^; LC, littermate control; myd, untreated *myd*; myd AAV, AAV*Large1* injected *myd* mice; IP, intraperitoneal; RO, retroorbital.

AAV-mediated gene transfer resulted in overexpression of *Large1* in *myd* mice, leading to hyperglycosylation of α-DG in the muscle sarcolemma (Figs. 2a, 2b) as previously observed with adenovirus expressing *Large1* (12). Western blotting of WGA-enriched muscle homogenates after AAV-mediated systemic transduction showed that α-DG was highly glycosylated with matriglycan and bound laminin (Fig. 2b). Furthermore, restoring *Large1* expression in *myd* mice rescued α-DG-ligand binding (Fig. 2c), explaining the observation that EDL muscles were protected from ECC-induced damage.

The improved phenotype in muscle size and function observed in *myd* mice upon AAV-mediated gene transfer of *Large1* prompted us to determine if this treatment also improved skeletal muscle pathology. As expected, histological analysis of skeletal muscle in untreated *myd* mice showed marked variation in fiber size, numerous fibers with central nuclei, adipose tissue infiltration, and fibrosis (Fig. 2d). Immunofluorescence analysis also showed a lack of matriglycan on α-DG in untreated *myd* mice (Fig. 2d). In contrast, quantification of matriglycan-positive α-DG fibers revealed that almost all fibers expressed functional α-DG in *myd* mice after injection with AAV*Large1* (Fig. 2e). Furthermore, progression of fibrous tissue deposition was halted in treated mice when compared to untreated mice (Fig. 2f). In the absence of *Large1* expression, mice exhibited muscle wasting (Figs. S5c-f), associated with muscle fiber atrophy as revealed by an overall decrease in fiber diameter in untreated *myd* quadriceps femoris muscles (Fig. 2g), with a higher proportion of fiber diameters of 10–20 μm (Fig. 2h). In contrast, AAV-mediated overexpression of *Large1* increased overall muscle fiber diameter, as demonstrated by a greater percentage of fibers having a larger fiber diameter (Figs. 2g, 2h).

Respiratory function is impaired in *myd* mice (28-30), as demonstrated by a marked reduction in tidal volume (TV) and minute volume (MV) compared to WT mice or littermate controls (Figs. 3a, 3b). Moreover, *myd* mice displayed lower TV and MV normalized to body weight (TV/BW and MV/BW) than WT mice or littermate controls (Figs. 3c, 3d). A respiratory function test revealed that *myd* mice treated with AAV*Large1* had a higher TV and MV than untreated mice (Figs. 3a and b), indicating that treated mice exhibit reduced forced breathing compared to untreated mice. In addition, TV/BW was moderately increased in treated mice verses untreated mice, whereas MV/BW was unaffected (Figs. 3c, 3d), demonstrating that AAV-mediated overexpression of *Large1* ameliorated defects in respiratory muscle function.

**Figure 3.**
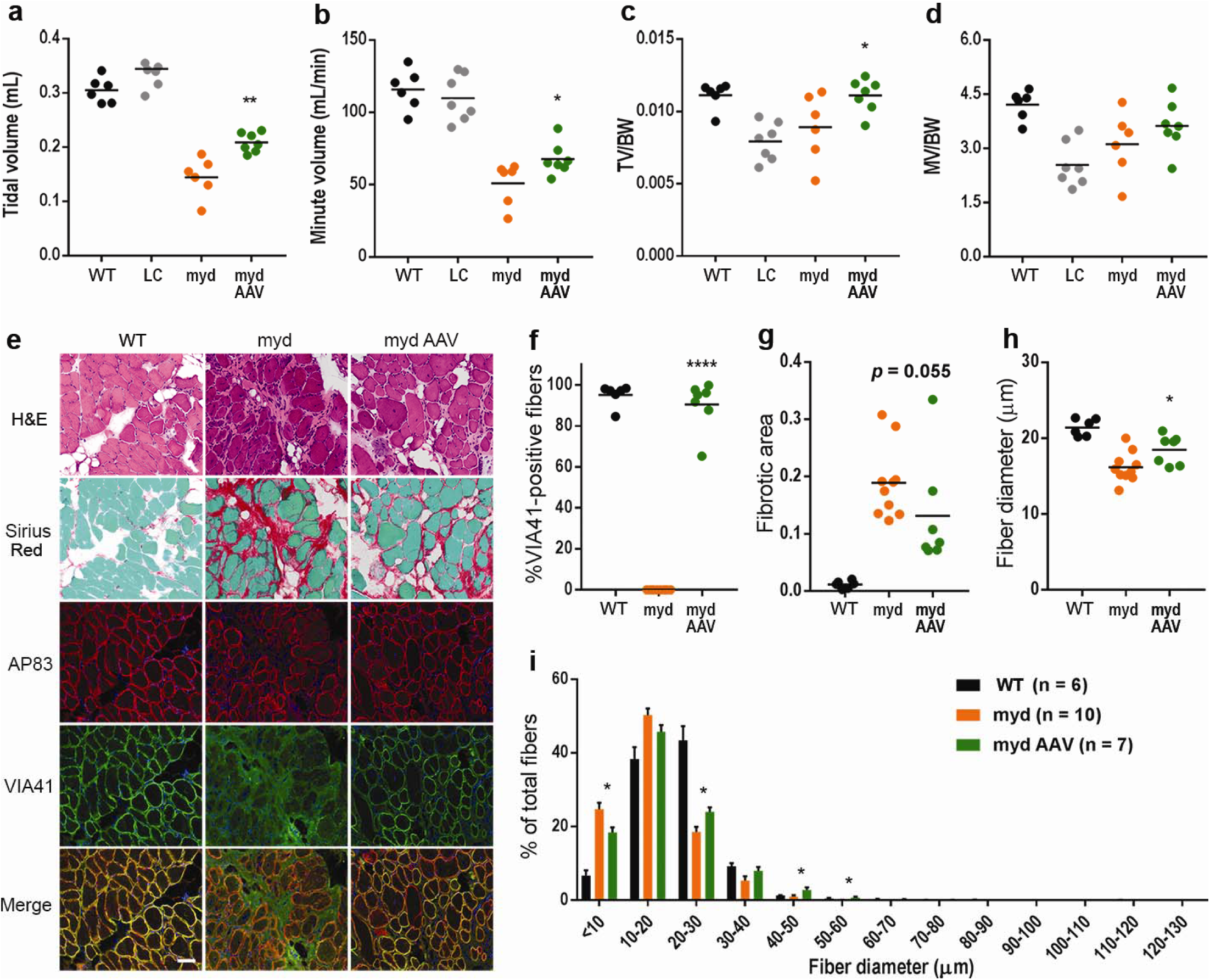
*Large1* gene transfer improves respiratory function and diaphragm pathology. Whole body plethysmography for: **a**, Tidal volume, TV; **b**, Minute volume, MV; **c**, Tidal volume normalized to bodyweight, TV/BW; **d**, Minute volume normalized to weight, MV/BW. **e**, Representative cryosections. Sections stained with H&E or Sirius Red & Fast Green, or used for immunofluorescence: β-DG (AP83) and matriglycan-positive α-DG (VIA41). Scale bar: 50μm. **f-i**, Quantitative analysis of sections in e. **f**, VIA41-positive fibers. **g**, Connective tissue deposition. **h**, Average Feret’s diameter of fibers. **i**, Percentage of fibers of indicated diameter. Symbols, individual mice; bars, means ± SEM. WT, C57BL/6J; myd, untreated *myd*; myd AAV, AAV*Large1* injected *myd* mice; LC, littermate control (*Large*^*+/+*^ or *Large*^+/-^).

We next investigated whether systemic gene transfer improves diaphragm pathology in *myd* mice. In untreated mice, fibrous tissue infiltration was prominent, and no muscle fibers expressed glycosylated α-DG (Figs. 3e, 3f). In contrast, treatment with AAV*Large1* restored expression of glycosylated α-DG to similar levels observed in WT mice (Fig. 3f). Progression of connective tissue deposition was also prevented by AAV-mediated gene transfer of *Large1* relative to untreated mice (Fig. 3g). In addition, the overall fiber diameter of diaphragm muscles was increased (Fig. 3h), and an increased percentage of muscle fibers had a larger fiber diameter upon overexpression of *Large1* (Fig. 3i). Thus, AAV-mediated gene transfer of *Large1* caused muscle fibers in the diaphragm to increase in diameter, thus improving respiratory function.

We further investigated whether systemic gene delivery using AAV restores matriglycan on α-DG in cardiomyocytes in *myd* mice. No obvious pathology was observed in the absence of *Large1* (Fig. S6a). Immunofluorescence analysis revealed that expression of glycosylated α-DG was remarkably restored in hearts of *myd* mice treated with AAV*Large1* and was likely due to restored expression of the *Large1* gene (Figs. S6a, 6b). Of note, the subendocardial portion was less transduced (Fig. S6a) indicating AAV transduction is less efficient in this region. No difference in connective tissue infiltration was observed between untreated and treated mice, although two mice that were treated with AAV*Large1* had focal fibrous tissue deposition (Fig. S6c). Therefore, AAV-mediated gene delivery of *Large1* restores glycosylated α-DG in cardiomyocytes, although the effects of this on cardiomyocyte function are difficult to discern in the *myd* mouse given the lack of phenotype in the heart in our *myd* mouse colony.

Skeletal muscle metabolism is a critical regulator of systemic metabolism that becomes abnormal in muscular dystrophies (31-34). The plasma metabolome is a useful read-out of whole-body metabolism that reflects changes in muscle metabolism resulting from both altered basal nutrient utilization and locomotive activity. To determine the efficacy of AAV*Large1* gene transfer in normalizing whole-body metabolism, we performed targeted metabolomic profiling on plasma from *myd* mice with advanced disease. The plasma metabolome was abnormal in *myd* mice, with significant changes observed in 21 of 78 measured plasma metabolites, 10 of which were normalized upon AAV*Large1* gene transfer (Table S1). Notably, glycolytic (Fig 4a) and TCA cycle (Fig 4b) intermediates were enriched among altered metabolites, especially those restored by AAV*Large1. myd* mice exhibited increased early and mid-glycolytic intermediates and decreased lactate, suggesting altered coupling of early and late glycolysis (Fig 4a). In parallel, TCA cycle intermediates malate, fumarate, and α-ketoglutarate were decreased, consistent with decreased TCA cycle amino acid influx through α-ketoglutarate in the absence of *Large1* (Fig 4b). Overall, the metabolomic profiling of plasma showed that systemic metabolism normalized following treatment with AAV*Large1*.

**Figure 4.**
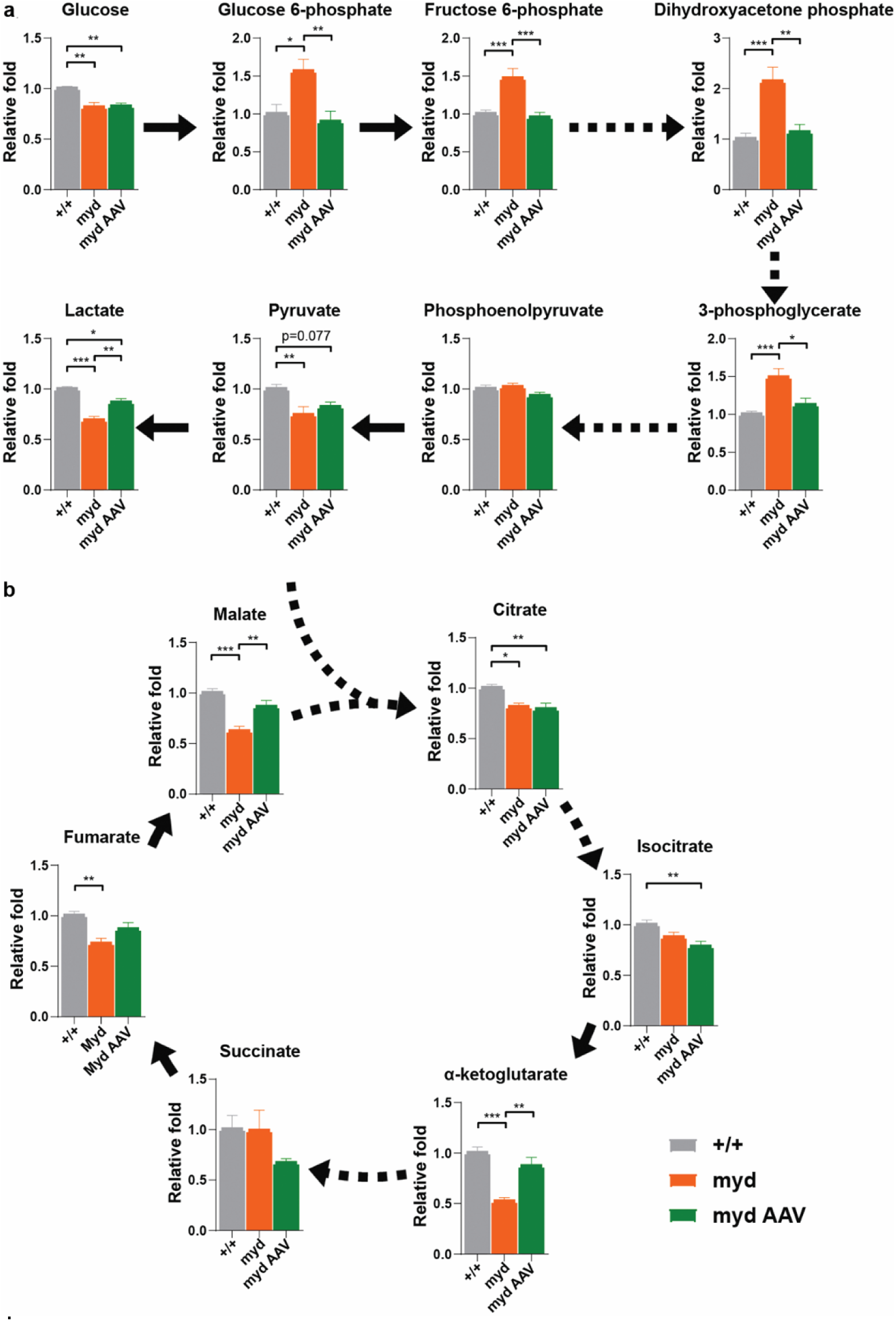
*Large1* is required for normal glycolytic and TCA cycle plasma profiles. Relative fold change of plasma: **a**, glycolytic and **b**, intermediate TCA cycle metabolites from *Large*^*+/+*^ (+/+, grey bars, n=11), untreated *myd* (myd, orange bars, n=7), and *myd* mice treated with AAV*Large1* (myd AAV, green bars, n=6), solid and dashed arrows represent single and multiple metabolic reactions, respectively, *p<0.05, **p<0.01, ***p<0.001. Analysis by ordinary one-way ANOVA followed by post hoc Tukey’s multiple comparison test. Data presented as mean ± SEM.

## DISCUSSION

Current therapies available to muscular dystrophy patients are limited and neither restore normal skeletal muscle function nor improve survival. Gene therapy is an exciting therapeutic approach for the treatment of muscular dystrophy as it enables the delivery of a functional copy of a gene or repairs the mutated locus. Preclinical studies using AAV-mediated gene transfer or gene editing to treat various mouse models of muscular dystrophy at a young age have been very successful in preventing skeletal muscle pathophysiology (7-15). However, the use of gene transfer has been difficult to study when applied as a treatment in the late stages, after the onset of severe muscle pathophysiology. Yet, these “restoration” studies are particularly important since many patients with muscular dystrophy have skeletal muscle degeneration before the onset of clinical symptoms and are often not diagnosed or treated until after these symptoms arise. Thus, we seek to understand the ability of gene therapy to repair severely damaged skeletal muscle and improve survival.

Our previous work with *Large1* gene transfer using adenovirus required direct muscle injections into young *myd* mice and showed localized prevention of dystrophic pathology (12). In addition, *Large1* gene transfer into healthy muscle did not cause abnormalities even though *Large1* was highly overexpressed and α-DG was hyperglycosylated (12). Here, in order to test if severely damaged skeletal muscle can be repaired and overall lifespan improved, we performed a comprehensive study using systemic AAV-mediated gene transfer of *Large1* in *myd* mice with advanced disease. Since α-DG and *Large1* are widely expressed, we used the CMV promoter to drive *Large1* expression and ensure α-DG glycosylation was restored. To evaluate the effectiveness of AAV-mediated gene transfer of *Large1*, we used various therapeutic readouts, including body weight, survival, muscle and respiratory function, muscle physiology, histological evaluation, plasma metabolomics, and biochemical analysis of α-DG.

We first used non-invasive techniques to monitor the effectiveness of systemic treatment with AAV*Large1* to restore muscle function in older *myd* mice. Both locomotor and rearing activities were restored, and mice were able to maintain motor performance. AAV*Large1-*treated *myd* mice also showed a significant increase in forelimb grip strength. Moreover, AAV-mediated expression of *Large1* ameliorated defects in respiratory muscle function and normalized systemic metabolism, as determined by metabolomic profiling on plasma. Although the precise mechanisms underlying these metabolomic changes must still be determined, these results identify plasma metabolic signatures that are associated with the absence of *Large1* in *myd* mice, and that are restored upon AAV*Large1* gene transfer. Consequently, future studies should avail themselves to plasma metabolomics, as it is a non-invasive, powerful, multivariate method to surveil the efficacy of gene therapy.

In addition, untreated *myd* mice displayed a significant decline in body weight that progressed until euthanasia was required, whereas systemic gene transfer of *Large1* resulted in a mild gain in body weight. Perhaps the most remarkable was the ability of AAV*Large1* to significantly extend the survival of older *myd* mice. Strikingly, all but one mouse injected with AAV*Large1* lived to >65 weeks of age, whereas only 50% of untreated *myd* mice survived to 45 weeks of age. Thus, results from our non-invasive assays provide evidence that α-DG function is restored following systemic treatment with AAV*Large1*, and that this has a significant impact on survival.

To further support our findings from non-invasive assays, we performed invasive analysis of muscle following euthanasia of mice treated with AAV*Large1*. Western blotting analysis and quantitative solid phase laminin binding assays demonstrated that AAV*Large1* was able to restore fully functional glycosylation of α-DG. Systemic transduction with AAV*Large1* in *myd* mice resulted in a significant increase in isometric tetanic force (F_o_) and specific isometric tetanic force (F_o_/CSA) of EDL muscles compared to untreated mice, demonstrating that muscle contractile properties can be improved, even at advanced stages of disease. Additionally, EDL muscles in AAV*Large1-*treated mice tolerated a sequence of eccentric contractions in a manner that was largely similar to that of WT mice. Collectively, our histological, biochemical, and physiological analysis of muscle show that glycosylation of α-DG is restored, and muscle function improved in *myd* mice treated with AAV*Large1*.

Individuals with muscular dystrophy are commonly diagnosed after the onset of clinical symptoms, yet current treatments are unable to restore the function of degenerated muscles. We show that AAV*Large1* gene transfer in older *myd* mice with severe muscular dystrophy attenuates skeletal muscle degeneration, improves motor and respiratory function, and normalizes systemic metabolism, which collectively extends survival of *myd* mice. Furthermore, our results demonstrate that skeletal muscle has remarkable plasticity, suggesting that gene therapy that is aimed at rescuing deficits in gene function harbors the potential to effectively restore muscle function. Thus, this study provides proof-of-concept that functional and pathological features of muscular dystrophy are reversible, even in advanced stages of disease.

## METHODS

### Experimental design

We set out to evaluate the therapeutic effect of treating established α-dystroglycanopathy; therefore, we targeted aged, severely affected *Large*^*myd*^/*Large*^*myd*^ (*myd*) mice as they exhibit end-stage muscle pathophysiology. The *myd* mouse is a natural model of glycosylation-deficient muscular dystrophy (14), with 65% of animals dying before 40 weeks of age (7–9). Previous work involving *Large1* gene transfer with adenovirus used direct muscle injections into young *myd* mice and showed localized prevention of dystrophic pathology (12). Successful systemic gene transfer using 10^12^-10^13^ vg/mouse via intraperitoneal or intravenous injections has previously been established for serotype 9, especially in skeletal muscle and heart (8-13). Therefore, we choose to use the AAV2/9 vector with a CMV promoter to systemically administer *Large1* to *myd* mice. Thirty-five surviving *myd* mice that were 34 weeks or older were randomly assigned to untreated or treated groups. In the treated group, a total of 14 mice were injected once intraperitoneally (n = 3) or once intravenously (n = 11) with AAV2/9CMV*Large1* (AAV*Large1*) at a vector dose of 4.35 × 10^12^ vg/mouse. Twenty-one mice were left untreated. Therapeutic readouts were body weight, survival, muscle and respiratory function, muscle physiology, histological evaluation, plasma metabolomics, and biochemical analysis of α-DG (see Fig. 1a for experimental outline). The mice were weighed weekly and locomotor activity was assessed every four weeks. The respiratory function test was performed just before untreated mice were euthanized or when AAV-treated mice became 60 weeks old.

### Animal Care and Criterion for euthanasia

Mice were maintained in a barrier-free, specific pathogen-free grade facility and had access to normal chow and water *ad libitum*. All animals were manipulated in biosafety cabinets and change stations using aseptic procedures. The mice were maintained in a climate-controlled environment at 25°C on a 12/12 hour light/dark cycle. All animal protocols were approved by the University of Iowa Animal Care and Use Committee (IACUC). MYD/Le-*Os*+/+*Large*^*myd*^/J mice (JAX stock #000300) were maintained on a C57BL/6J (WT) background and colony maintenance was carried out in the laboratory by mating +/*myd* males to +/*myd* females. *Myd* mice and control littermate mice (*+*/+ or +/*myd* mice) were identified via PCR.

The genetic background of our *myd/myd* (MYD/Le-*Os+/+Large*^*myd*^/J) mouse line (Jax#000300) was tested at Transnetyx (Cordova, TN) with an array-based platform using over 10,000 SNP markers, which showed the line has a 92.5% C57BL/6 genetic background. Animal care, ethical usage, and procedures were approved and performed in accordance with the standards set for the by the National Institutes of Health and IACUC. Of the 161 offspring from heterozygous +/*myd* matings, only 18 (11.2%) were homozygotes (*myd*/*myd*). Theoretically, there is a 25 % chance that a homozygote is produced, suggesting some homozygotes could be embryonic lethal. Surviving *myd* mice older than 35 weeks of age required gruel feeding in the cage, as they were too weak to reach food pellets suspended overhead. Due to the progressive deterioration in motor and respiratory function with age, *myd* mice were euthanized due to an inability to feed, hind limb paralysis, reduced body score, or respiratory distress, as dictated in our IACUC protocol.

### AAV vector production and AAV injection

The sequence encoding mouse *like-acetylglucosaminyltransferase-1* (*Large1*) was synthesized (Genscript, Piscataway, NJ) and cloned into the AAV backbone under the transcriptional control of the ubiquitous CMV promoter. The AAV2/9 vector contains the genome of serotype 2 packaged in the capsid from serotype 9 and was selected due to its ability to improve muscle transduction efficiency as well as alter tropism. The vector AAV2/9CMV*Large1* was generated by the University of Iowa Viral Vector Core Facility. For adult mice, 100 microliters (4.35 × 10^12^ vg) of the vector solution was administered once intraperitoneally or intravenously via the retro orbital (RO) sinus.

### Locomotor activity

Locomotor activity was measured using an open-field Digiscan apparatus (Omnitech Electronics, Columbus, OH). The mice were acclimated to the open-field apparatus for three days prior to the first trial. Total walking distance and rearing behavior (vertical activity) were recorded every 10 minutes for one hour when the mice were in active phase. Activity data were collected for two consecutive days and the higher of the two values for each parameter was used for analysis.

### Forelimb grip strength test

The forelimb grip strength test was performed when mice were eight weeks of age in the preventive study and 60 weeks of age in the therapeutic study. A mouse grip strength meter (Columbus Instruments, Columbus, OH) was mounted horizontally, with a nonflexible grid connected to the force transducer, which shows the highest force applied by the mouse on the grid during the pull. The mouse was allowed to grasp the grid with its front paws and then pulled away from the grid so that its grasp was broken. The gram force was recorded per pull, but measures were rejected in which only one forepaw or the hind limbs were used, and in which the mouse turned during the pull. A total of 15 pulls (five series of three pulls in a row) were performed, with a resting period between series to allow the mouse to recover and avoid habit formation. The three highest values out of the 15 values collected were used to determine the maximum grip strength, which was normalized for body weight.

### Plethysmography

Respiratory function was tested in unrestrained mice using a whole-body plethysmograph (Buxco Respiratory Products, Data Sciences International, St. Paul, MN). The mouse was weighed and placed into the chamber for 15 minutes to acclimate and respiratory flow data was recorded for five minutes using FinePointe software (Data Sciences International, St. Paul, MN). The measurements were done for two consecutive days. For data analysis, average values for tidal volume (TV), minute volume (MV), TV normalized by body weight (TV/BW), and MV normalized by body weight (MV/BW) were used.

### Measurement of *in vitro* muscle function

To compare the contractile properties of muscles, EDL muscles were surgically removed and analyzed as described previously (35, 36), with minor modifications. The muscle was immediately placed in a bath containing a buffered physiological salt solution (composition in mM: NaCl, 137; KCl, 5; CaCl_2_, 2; MgSO_4_, 1; NaH_2_PO_4_, 1; NaHCO_3_, 24; glucose, 11). The bath was maintained at 25°C, and the solution was bubbled with 95% O_2_ and 5% CO_2_ to stabilize the pH at 7.4. The proximal tendon was clamped to a post and the distal tendon was tied to a dual mode servomotor (Model 305C; Aurora Scientific, Aurora, ON, Canada). Optimal current and whole muscle length (L_o_) were determined by monitoring isometric twitch force. Optimal frequency and maximal isometric tetanic force (F_o_) were also determined. The muscle was then subjected to an eccentric contraction (ECC) protocol consisting of five eccentric contractions (ECCs) separated by three-minute intervals. A fiber length (L_f_)-to-L_o_ ratio of 0.45 was used to calculate L_f_. Each ECC consisted of an initial 100 ms isometric contraction at optimal frequency immediately followed by a stretch of L_o_ to 30% of L_f_ beyond L_o_ at a velocity of one L_f_/s at optimal frequency. The muscle was then passively returned to L_o_ at the same velocity. At three, 15, 30, 45, and 60 minutes after the ECC protocol, isometric tetanic force was measured, and force deficit was calculated as the decrease in isometric tetanic force post-ECC as a percentage of F_o_. After the analysis of the contractile properties, the muscle was weighed. The cross-sectional area (CSA) of muscle was determined by dividing the muscle mass by the product of L_f_ and the density of mammalian skeletal muscle (1.06 g/cm^3^). The specific force was determined by dividing F_o_ by the CSA (mN/mm^2^).

### Tissue collection and histological evaluation

Mice were euthanized by cervical dislocation and tissue samples were obtained. Quadriceps, gastrocnemius, tibialis anterior, and soleus muscles were weighed prior to processing. Quadriceps and gastrocnemius muscles from the right side of the animal, half the diaphragm, and half the heart were snap frozen in Optimal Cutting Media (OCT), submerged in isopentane cooled in liquid nitrogen, and stored at -80 °C for further analysis. The other skeletal muscles were frozen in liquid nitrogen and stored at -80 °C for biochemical analysis. Cryosections of quadricep muscle, diaphragm, and heart were cut at a thickness of 7 μm and stained with hematoxylin and eosin (H&E) and Sirius red and Fast Green. For Sirius red staining, sections were fixed in 10% neutral buffered formalin (Thermo Fisher Scientific) for five minutes, followed by Sirius red staining consisting of a 30-minute incubation in 0.1% Sirius red F3B (1A 280; Chroma Gesellschaft, Germany) in saturated picric acid (Sigma-Aldrich, St Louis, MO), and several incubations with ethyl alcohol and xylene. Whole digital images of H&E- and Sirius red-stained sections were taken by a VS120-S5-FL Olympus slide scanner microscope (Olympus Corporation, Tokyo, Japan). To quantify fibrous tissue infiltration in Sirius red-stained muscle and diaphragm sections, four fields were randomly selected from a whole section using OlyVIA ver.2.9 (Olympus) and the captured images were processed using Image J (NIH) with additional threshold color plug-ins to process jpeg images. Pixels corresponding to the area stained in red were normalized to the total pixel area of the image, and the four values were expressed as a percent of the fibrotic area and were averaged for each section.

### Immunohistochemistry

For morphometric analyses, transverse sections of the sarcolemma of muscle, diaphragm, and heart were stained with a rabbit polyclonal anti-caveolin three antibody (ab2912; Abcam, 1:100 dilution) and a mouse monoclonal antibody to glycoepitopes on the sugar chain of α-DG (VIA41; 1:10 dilution) overnight at 4°C, followed by staining with Alexa Fluor®-conjugated goat IgG against rabbit IgG and goat IgG against mouse IgG_1_ (Invitrogen, 1:400 dilution), respectively, for 40 minutes. The sections were counterstained with DAPI (Invitrogen) and whole sections were imaged with a VS120-S5-FL Olympus slide scanner microscope. Feret’s diameter of muscle fibers were measured with VS-DESKTOP software (Olympus). To quantify fibers expressing glycosylated α-dystroglycan, the percent of VIA41-positive fibers was determined by dividing the VIA41-positive fiber count by the anti-caveolin 3-positive fiber count. To quantify fibers with centrally located nuclei, three fields were randomly selected from a whole section and were double stained with anti-caveolin 3 antibody and DAPI. Total and centrally nucleated fibers (CNFs) were manually counted and %CNF was determined by dividing CNF by total fiber count per each captured image. The three %CNF values were averaged and compared. The sarcolemma of muscle and diaphragm sections were also stained with a rabbit monoclonal antibody to β-DG (AP83; 1:25 dilution) and VIA41. IIH6 and VIA41 antibodies are monoclonal antibodies to the glycoepitope of α-DG (20, 21), and AP83 is a polyclonal antibody to the Ce-terminus of β-DG (19), all of which have been described previously.

### Glycoprotein enrichment and Western blot analysis

Half of the quadricep muscle was solubilized in 1 mL Tris-buffered saline (TBS) containing 1% Triton X-100 and protease inhibitors. The solubilized fraction was incubated with 200 microliters of WGA-agarose bead slurry (Vector Laboratories, Burlingame, CA) overnight at 4°C. Pellets formed from the beads and were washed three times in 1 mL TBS containing 0.1% Triton X-100 (15). The beads were then either directly mixed with SDS-polyacrylamide gel electrophoresis (PAGE) loading buffer (for western blotting, ligand overlay) or eluted with 1 mL TBS containing 0.1% Triton X-100 and 300 mM *N*-acetyl-glucosamine (for solid-phase binding assay). Proteins were separated by 3–15% SDS-PAGE and transferred to polyvinylidene fluoride-FL (PVDF-FL, Millipore Sigma) membranes. The membranes were incubated with a sheep polyclonal antibody to human DG (AF6868; R&D Systems, 1:100 dilution) and a mouse monoclonal antibody to a glycoepitope on the sugar chain of α-DG (IIH6; 1:100 dilution) followed by IRDye® 800CW dye-conjugated goat anti-sheep IgG (LI-COR, 926-32214) and goat anti-mouse IgM (LI-COR, 926-32280), respectively.

### Ligand overlay assay

Ligand overlay assays were performed on PVDF-FL membranes using mouse Engelbreth– Holm–Swarm (EHS) laminin (ThermoFisher, 23017015). Briefly, PVDF-FL membranes were blocked in laminin binding buffer (LBB: 10 mM triethanolamine, 140 mM NaCl, 1 mM MgCl_2_, 1 mM CaCl_2_, pH 7.6) containing 5% milk followed by incubation with laminin overnight at 4°C in LBB containing 3% bovine serum albumin (BSA). Membranes were washed and incubated with anti-laminin antibody (L9393; Sigma-Aldrich, 1:100 dilution) followed by IRDye® 800CW dye-conjugated donkey anti-rabbit IgG (LI-COR, 926-32213).

### Solid-phase assay

A solid-phase laminin binding assay was performed as described previously (14). Briefly, WGA eluates were diluted 1:50 in TBS and coated on polystyrene ELISA microplates (Costar 3590) overnight at 4°C. Plates were washed in LBB and blocked for two hours in 3% BSA/LBB at RT. Mouse EHS laminin was diluted in 1% BSA/LBB and applied for one hour at RT. The wells were washed with 1% BSA/LBB and incubated for one hour with L9393 (1:5,000 dilution) in 3% BSA/LBB followed by incubation with HRP-conjugated anti-rabbit IgG (Invitrogen, 1:5,000 dilution) in 3% BSA/LBB for 30 minutes. Plates were developed with o-phenylenediamine dihydrochloride and H_2_O_2_, and reactions were stopped with 2 N H_2_SO_4_. Absorbance per well was read at 490 nm by a microplate reader.

### mRNA expression analysis

Quadricep muscle and heart cryosections were collected into tubes and stored at ™80°C until use. Total RNA was isolated using an RNeasy Mini kit (Qiagen, Venlo, Netherlands) and cDNA was generated using a QuantiTect Reverse Transcription Kit (Qiagen). For gene expression analysis, a QX100 droplet digital PCR (ddPCR) system (Bio-Rad, Pleasanton, CA) was used. The ddPCR reaction mixture consisted of 10 microliters of a 2× EvaGreen Supermix (Bio-Rad), 2 microliters of primers, and 5 microliters of cDNA sample in a final volume of 20 microliters. The entire reaction mixture was loaded into a disposable plastic cartridge (Bio-Rad) together with 70 microliters of droplet generation oil (Bio-Rad) and placed in the droplet generator (Bio-Rad). After processing, the droplets generated from each sample were transferred to a 96-well PCR plate (Eppendorf, Hamburg, Germany). PCR amplification was carried out on a T100 thermal cycler (Bio-Rad) using a thermal profile beginning at 95°C for five minutes, followed by 40 cycles of 95°C for 30 seconds and 60°C for 60 seconds, one cycle of 90°C for five minutes, and ending at 4°C. After amplification, the plate was loaded on the droplet reader (Bio-Rad) and the droplets from each well of the plate were read automatically at a rate of 32 wells per hour. ddPCR data were analyzed with QuantaSoft analysis software (Bio-Rad), and the quantification of the target molecule was presented as the number of copies per microliter of PCR mixture. The oligonucleotides used for amplification are listed in Supplementary Table S2.

### Targeted metabolomics

Forty microliters of plasma was vortexed with 720 microliters of ice cold 1:1 methanol/acetonitrile to extract metabolites and incubated for one hour at –20°C. Metabolite extracts were centrifuged for 10 minutes at 21,000 x g to pellet precipitated protein. Supernatants were transferred to sample vials and dried using a speed-vac. The resulting dried metabolite extracts were derivatized using methoxyamine hydrochloride (MOX) and N,O-Bis(trimethylsilyl)trifluoroacetamide (TMS) and examined by gas chromatography-mass spectrometry (GC-MS), as previously described (37, 38). Briefly, dried extracts were reconstituted in 30 microliters of 11.4 mg/ml MOX in anhydrous pyridine, vortexed for five minutes, and heated for one hour at 60°C. Next, 20 microliters of TMS was added to each sample, which were then vortexed for one minute and heated for 30 minutes at 60°C. Samples were immediately analyzed using GC-MS. GC separation was conducted on a Thermo Trace 1300 GC fitted with a TraceGold TG-5SilMS column. One microliter of derivatized sample was injected into the GC operating under the following conditions: split ratio = 20-1, split flow = 24 microliters/minute, purge flow = 5 ml/minute, carrier mode = Constant Flow, and carrier flow rate = 1.2 ml/minute. The GC oven temperature gradient was as follows: 80°C for 3 minutes, increasing at a rate of 20°C/minute to 280°C, and holding at a temperature at 280°C for 8 minutes. Metabolites were detected using a Thermo ISQ 7000 mass spectrometer operated from 3.90 to 21.00 minutes in EI mode (−70eV) using select ion monitoring (SIM). EI fragmented metabolites were identified by the m/z of fragments at unique chromatographic retention times corresponding to previously analyzed reference standards. Peak intensities were corrected using the NOREVA tool (39). Peak intensities from six unique experiments were normalized to total signal per sample and set relative to WT for final analysis.

### Statistical analysis

All data in the present study are shown as the means ± SEM unless otherwise indicated. The number of sampled units, n, upon which we reported statistics, is the single mouse for the *in vivo* experiments (one mouse is n = 1). GraphPad Prism 7 software was used for statistical analyses. The statistical analysis performed for each data set is indicated in the figure legends. P < 0.05 was considered significant. For survival, treated mice were compared with untreated mice by Kaplan-Meier log-rank test. For all figures, \**p* < 0.05, \*\**p* < 0.01, \*\*\**p* < 0.001, and \*\*\*\**p* < 0.0001 were used.

## END NOTES

## Acknowledgements

We thank Keith Garringer for technical assistance and the Viral Vector Core at the University of Iowa for generating the adeno-associated viral vector. ddPCR analyses presented herein were obtained at the Genomics Division of the Iowa Institute of Human Genetics, which is supported, in part, by the University of Iowa Carver College of Medicine. We are grateful to Drs. Jennifer Barr and Christine Blaumueller of the Scientific Editing and Research Communication Core at the University of Iowa Carver College of Medicine for critical reading of the manuscript. We are grateful to Amber Mower and Jaeda Harmon for assistance with administrative support.

## Authors’ contributions

T.Y. co-designed the project, carried out the experimental work, analyzed and interpreted the data, and co-wrote the manuscript. S.E.H., M.A.C., M.E.A., A.J.R., H.O., and A.D.P. conducted experimental work and analyzed and interpreted the data. D.V. generated essential reagents. E.B.T. analyzed and interpreted the data, co-wrote the manuscript, and supervised the research. Corresponding author K.P.C. co-designed the project, analyzed and interpreted the data, co-wrote the manuscript, and supervised the research.

## Author Information

The authors declare no competing financial interests. Correspondence and requests for materials should be addressed to K.P.C.

## Supplementary Materials

Figures S1-S6

Data Tables S1 and S2

