## Supplementary material for "*Large1* Gene Transfer in Older *myd* Mice with Severe Muscular Dystrophy Restores Muscle Function and Greatly Improves Survival": Supplmentary Figures

**This PDF file includes:**

Figures S1-S6

Data Tables S1 and S2

**SUPPLEMENTAL FIGURES**

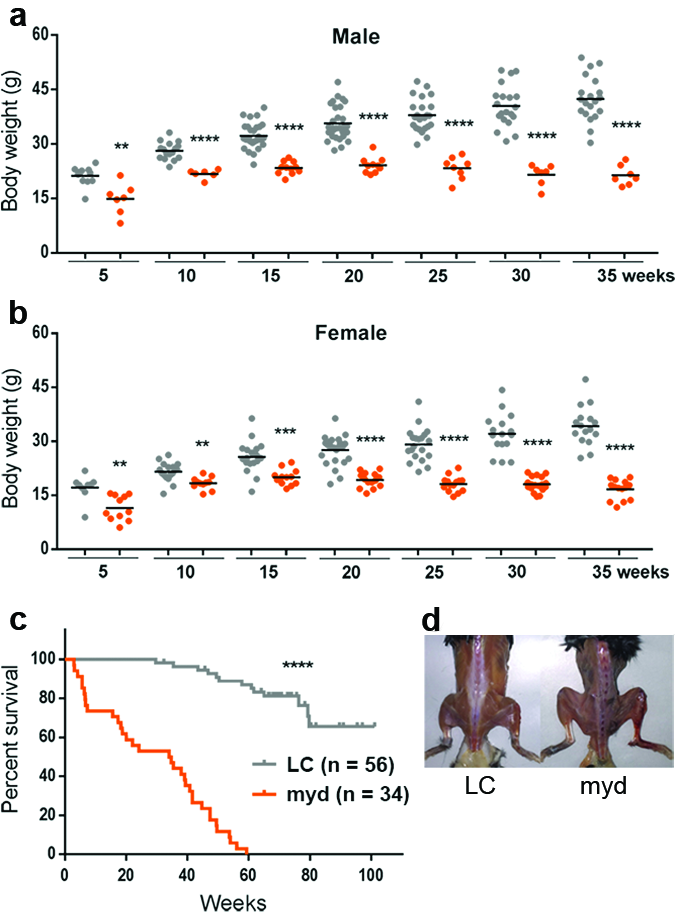

**Fig. S1 │** ***myd*** **mice show growth retardation, a shortened lifespan, and muscle wasting.**

**a,b,** Body weights of male (**a**; littermate control (LC): n=20, *myd*: n=7) and female (**b**; LC: n=16; *myd*: n=14) mice, which were weighed weekly, with weights compared at 5-week intervals. **c,** Percent survival of LC and *myd* mice. **d,** Image of a *myd* mouse depicting thoracic kyphosis and muscle wasting; 68.1-week-old LC mouse (left), 41.6-week-old *myd* mouse (right). For panels **a** and **b**, symbols represent individual mice, bars represent means. Littermate control (LC), gray symbols; *myd* (myd), orange symbols. * p<0.05; ** p<0.01, *** p<0.001; **** p<0.0001.

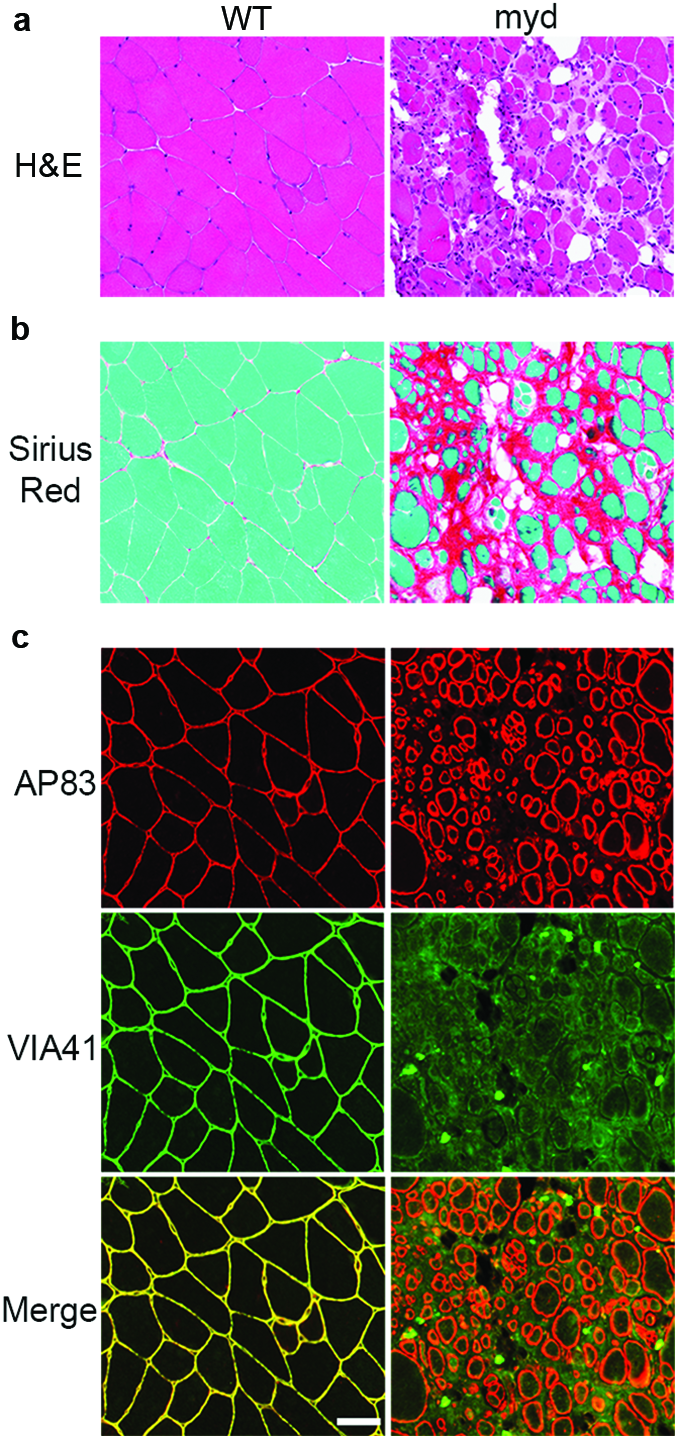

**Fig. S2 │ Quadricep muscles from aged *myd* mice exhibit severe dystrophic pathology and defects in glycosylation of α-DG.**

**a,b,** Quadricep muscle cryosections from a 69.0-week-old C57BL/6J (WT) mouse and a 41.6-week-old *myd* (myd) mouse stained with H&E or Sirius Red & Fast Green, or used immunofluorescence: β-DG (AP83); matriglycan-positive α-DG (VIA41). **c,** Immunofluorescence analysis of cryosections for β-DG (AP83) and glycosylated α-DG (VIA41). Scale bar represents 50 μm.

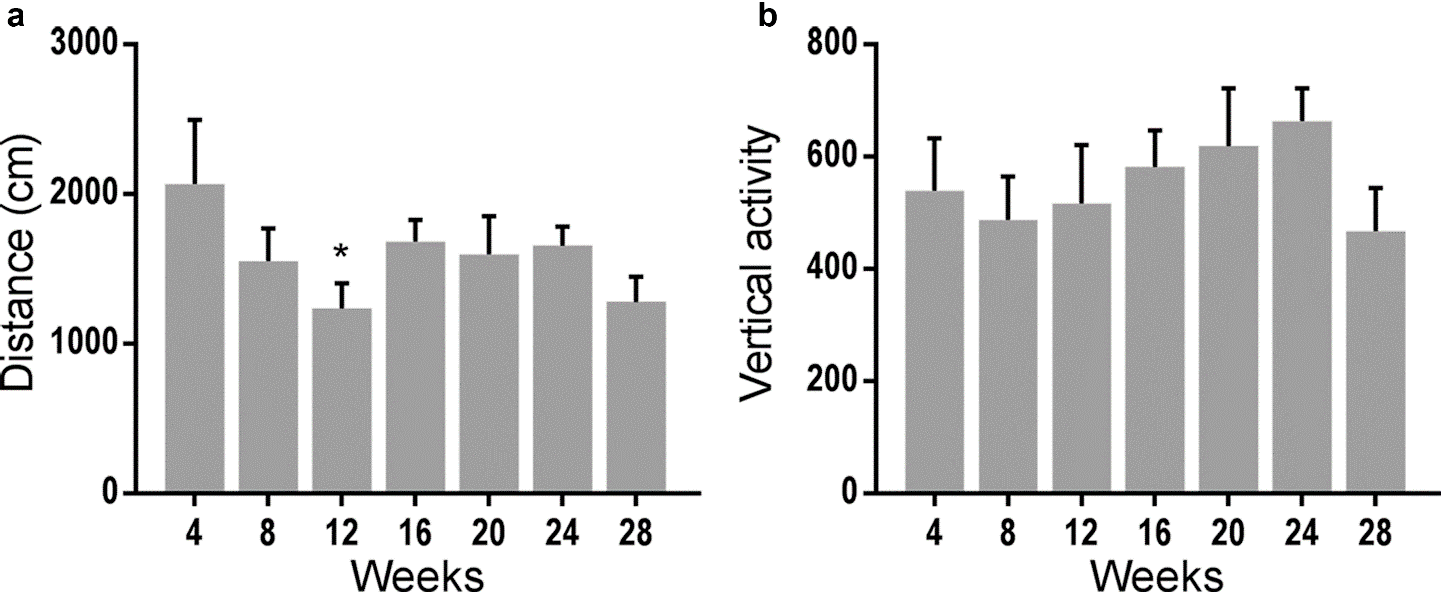

**Fig. S3 │Chronological evaluation of locomotion and rearing activity in C57BL/6J WT mice.**

**a,b,** Spontaneous locomotion (distance) and rearing (vertical activity) were measured every four weeks in C57Bl/6J WT mice. Paired t-tests were used to compare the parameters between baseline activity and that at 4, 8, 12, 16, 20, 24, and 28 weeks (n = 6). Bars represent mean ± SEM. * p<0.05.

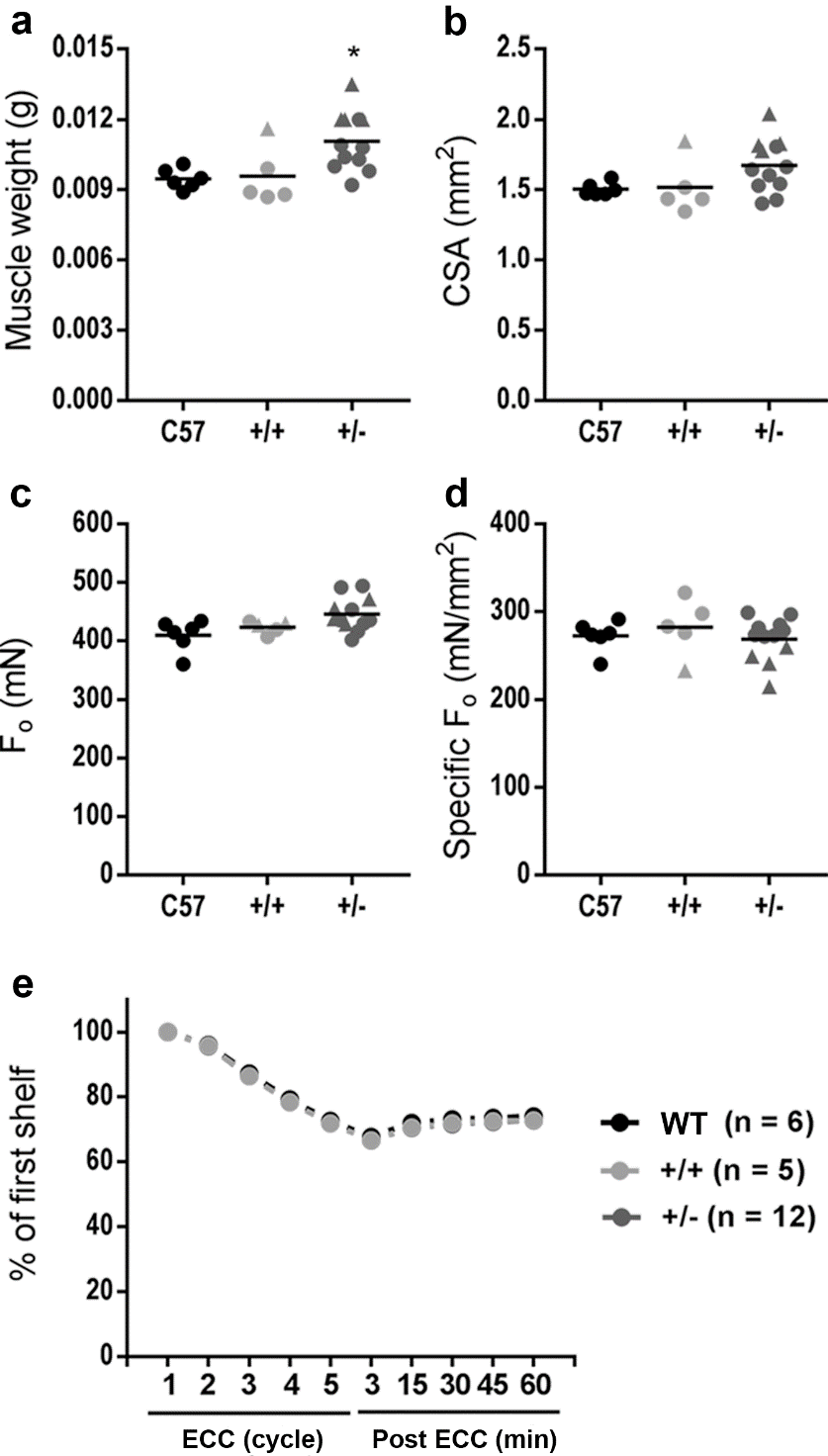

**Fig. S4 │Muscle size, contractile properties, and lengthening contraction-induced muscle damage is similar in EDL muscles from +/+ or +/*myd* mice.**

**a,** Muscle weight, **b** whole-muscle CSA, **c,** isometric tetanic force (F_o_), and **d,** specific isometric tetanic force (F_o_/CSA) were measured in C57Bl/6J (WT), +/+, and +/*myd* (+/-) mice. Unpaired t-tests were used to compare the parameters between +/+ and +/*myd* mice. **e,** Percentages of F_o_ at the 2^nd^, 3^rd^, 4^th^, or 5^th^ eccentric contraction (ECC) cycle, and at 3, 15, 30, 45, and 60 min after the eccentric contraction protocol (Post ECC) were calculated as a relative value to F_o_ at the 1^st^ ECC, in mice as in (a). Filled triangles, male; filled circles, female. C57, black symbols; +/+, light gray symbols; +/-, dark gray symbols. * p<0.05.

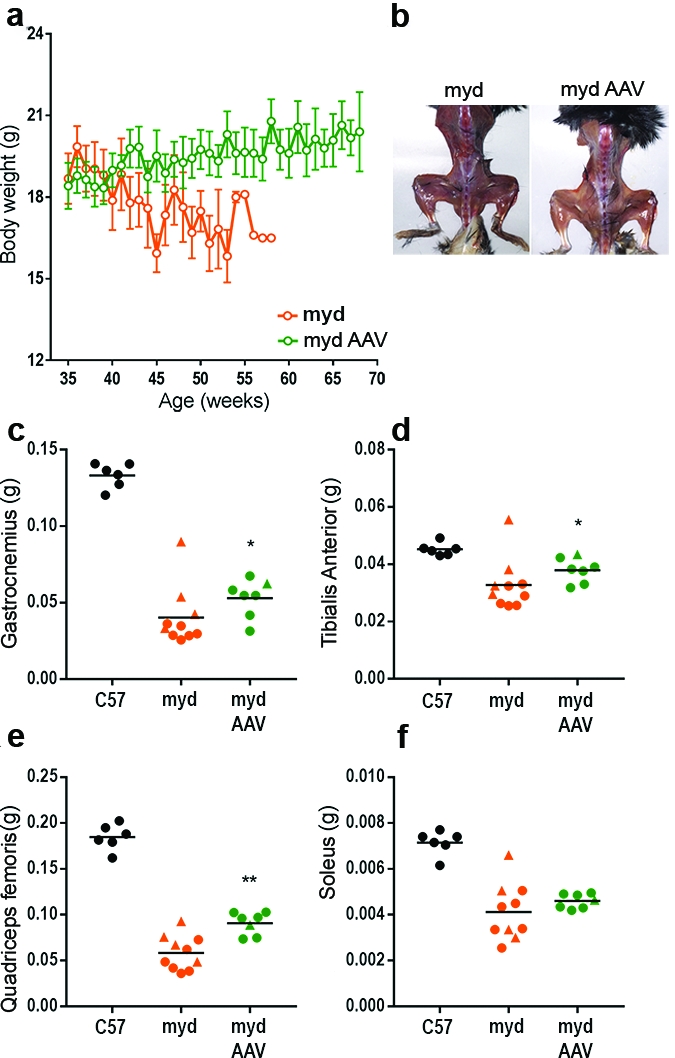

**Fig. S5 │AAV-mediated *Large1* gene transfer improves muscle mass in *myd* mice.**

**a,** Body weight of untreated (orange) or AAV*Large1*-treated (green) *myd* mice. Mice were treated with AAV*Large1* at >35 weeks and weighed weekly for at least 30 weeks. Linear regression analysis was performed, and both slopes were significantly non-zero: untreated, Y = -0.1125*X + 22.88; treated, Y = 0.05238*X + 16.85. **b,** Gross analysis of mice treated as in (**a**); 40.7-week-old untreated mouse (left), 68.1-week-old treated mouse (right). **c-f,** Weight of the gastrocnemius **c**, tibialis anterior **d**, quadriceps femoris **e**, and soleus **f,** muscles from C57Bl/6J (C57) mice, untreated (myd), or treated (myd AAV) *myd* mice as in (**a**). Average weights were compared between untreated and treated groups using Mann-Whitney or unpaired t-test. Symbols represent individual mice, bars represent mean ± SEM. Filled triangles, male; filled circles, female. C57, black symbols; myd, orange symbols; myd AAV, green symbols. * p<0.05; ** p<0.01.

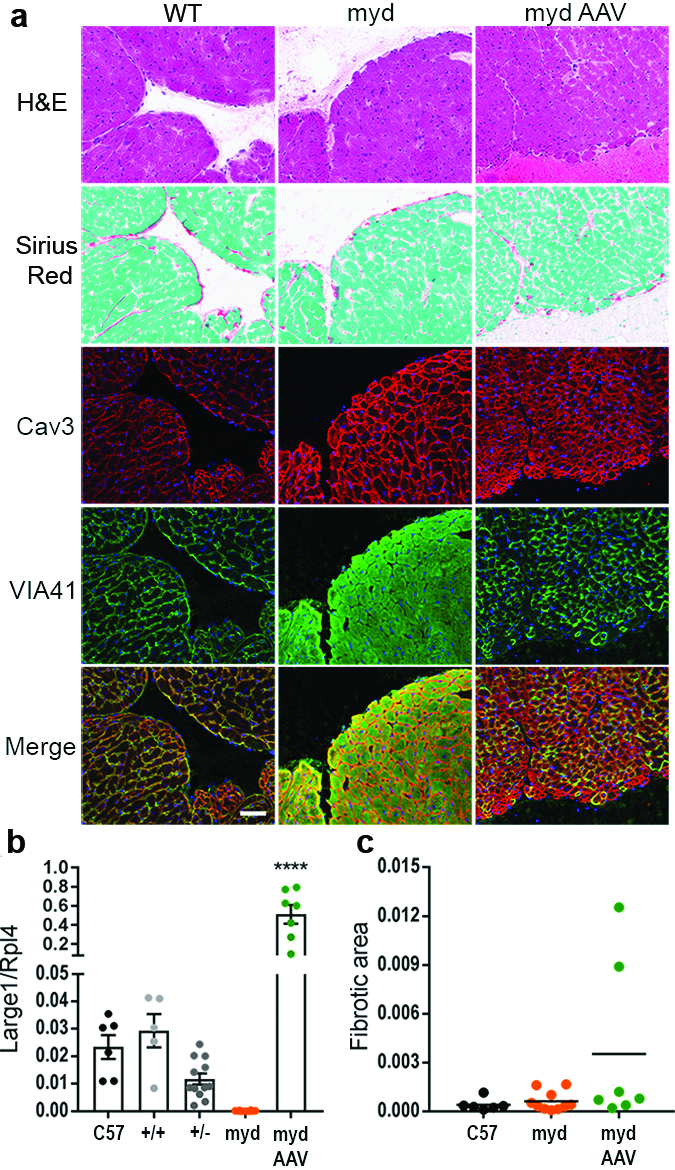

**Fig. S6 │Restoring Large1 expression improves muscle pathology, immunohistochemistry, and morphometric analysis** **of cardiac transverse sections.**

**a,** Representative cardiac transverse sections from a 68.1-week-old C57BL/6J (WT) mouse, a 56.1-week-old *myd* (myd) mouse, and a 69.3-week-old *myd* mouse treated with AAV*Large1* (myd AAV)*.* Sections stained with H&E or Sirius Red & Fast Green, or used immunofluorescence: β-DG (AP83); matriglycan-positive α-DG (VIA41). Scale bar represents 50μm. **b,** ddPCR analysis of *Large1* expression relative to *Rpl4* expression in cardiac muscle from C57Bl/6J mice (C57; n = 6), +/+ (n = 5) and +/*myd* (+/-; n = 12), untreated *myd* (myd; n = 10), and *myd* mice treated with AAV*Large1* (myd AAV; n = 7). Symbols represent individual mice, bars represent mean ± SEM. **c,** Connective tissue deposition to assess fibrosis, determined by Sirius Red & Fast Green staining in (**a**). Untreated and treated groups compared with unpaired t-test.

**SUPPLEMENTARY DATA TABLES**

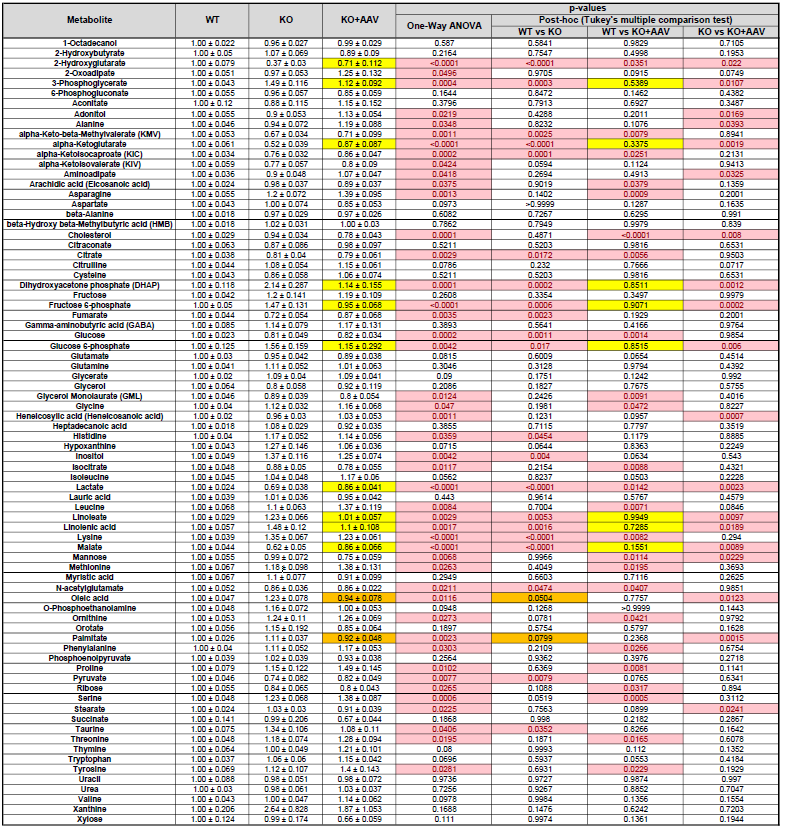
**Data Table 1 │Plasma Metabolomics**

Relative fold change of plasma metabolites in WT (+/+, n = 26), *myd* (myd, n = 20), and *myd* mice treated with AAV*Large1* (myd AAV, n = 22). Data are presented as mean ± SEM. Analysis by one-way ANOVA followed by Tukey’s post hoc multiple comparison test. p-valves from statistical analysis are reported; p-values <0.05 are shown in red text in a light-red filled cell. Yellow and gold highlighted cells emphasize the metabolites rescued by AAV*Large1* treatment.

**Data Table 2 │Primer design**

| Gene | Forward | Reverse | Amplicon size (bp) |
| --- | --- | --- | --- |
| *Large1* | acc tgc agt gcg agt aga c | cct gtt gcc ctt tga act tat gg | 224 |
| *Rpl4* | cca aga cta tgc gca gga at | tgt ctg cag tcc cct tct ct | 135 |
